# Natural genetic variation underlying tiller development in barley (*Hordeum vulgare* L)

**DOI:** 10.1101/730945

**Authors:** Allison M. Haaning, Kevin P. Smith, Gina L. Brown-Guedira, Shiaoman Chao, Priyanka Tyagi, Gary J. Muehlbauer

**Affiliations:** Department of Plant and Microbial Biology, University of Minnesota, Saint Paul, Minnesota, United States of America; Department of Agronomy and Plant Genetics, University of Minnesota, Saint Paul, Minnesota, United States of America; Plant Science Research, United States Department of Agriculture-Agricultural Research Service, Raleigh, North Carolina, United States of America; Department of Cereal Crops Research, Red River Valley Agricultural Research Center, United States Department of Agriculture-Agricultural Research Service, Raleigh, North Carolina, United States of America

**Keywords:** Genome-wide association, GWAS, barley, tiller, lateral branch

## Abstract

In barley (*Hordeum vulgare* L.), lateral branches called tillers contribute to grain yield and define shoot architecture, but genetic control of tiller number and developmental rate are not well characterized. The primary objectives of this work were to examine relationships between tiller number and other agronomic and morphological traits and identify natural genetic variation associated with tiller number and rate, and related traits. We grew 768 lines from the USDA National Small Grain Core Collection in the field and collected data over two years for tiller number and rate, and agronomic and morphological traits. Our results confirmed that spike row-type and days to heading are correlated with tiller number, and as much as 28% of tiller number variance is attributed to these traits. In addition, negative correlations between tiller number and leaf width and stem diameter were observed, indicating trade-offs between tiller development and other vegetative growth. Thirty-three quantitative trait loci (QTL) were associated with tiller number or rate. Of these, 40% overlapped QTL associated with days to heading and 22% overlapped QTL associated with spike row-type, further supporting that tiller development is influenced by these traits. Despite this, some QTL associated with tiller number or rate, including the major QTL on chromosome 3H, were not associated with any other traits, suggesting that tiller number can be modified independently of other important agronomic traits. These results enhance our knowledge of the genetic control of tiller development in barley, which is important for optimizing tiller number and rate for yield improvement.

## INTRODUCTION

Grasses form modified lateral branches called tillers that develop from axillary meristems (AXM) located in leaf axils near the base of the plant. Barley shoot architecture is largely defined by the number and vigor of tillers, which have the capacity, like the main shoot, to form grain-bearing inflorescences called spikes that contribute to grain yield (Cannell 1969). However, merely increasing tiller number may not increase grain yield because it has been associated with decreased seed number and seed weight, and increased lodging (Stoskopf and Reinbergs 1966; Simmons *et al.* 1982; Benbelkacem *et al.* 1984). Furthermore, tiller number is a complex trait influenced by photoperiod sensitivity, spike row-type, and environmental variables, including water and nitrogen availability and planting density (Turner *et al.* 2005; Alqudah and Schnurbusch 2014; Liller *et al.* 2015; Alqudah *et al.* 2016). Therefore, a more comprehensive understanding of the genetic basis of shoot architecture and relationships with other agronomic traits is important for altering barley shoot architecture for increased grain yield.

Tiller development (tillering) in barley has been characterized in several high and low tillering mutants, and five genes regulating tillering have been isolated and characterized to date. *LOW NUMBER OF TILLERS 1* (*LNT1*) encodes a BEL-like homeodomain transcription factor homologous to *Arabidopsis BELLRINGER* (*BLR*) and mutations in *LNT1* result in reduced tiller number (Dabbert *et al.* 2010). *UNICULME4* (*CUL4*) is homologous to *Arabidopsis BLADE-ON-PETIOLE* (*BOP*) genes and encodes a protein with a BROAD COMPLEX, TRAMTRACK, BRIC-À-BRAC (BTB)-ankyrin domain; and *cul4* mutants produce very few primary tillers and no secondary tillers (Tavakol *et al.* 2015). The *eligulum-a* (*eli-a*) mutant, was identified as a suppressor of the *uniculm2* (*cul2*) mutant phenotype (Okagaki *et al.* 2018). Typically, *cul2* mutants do not produce any tillers, but when combined with *eli-a* alleles, they develop at least one tiller. *ELI-A* encodes a conserved protein that may be a transposon, and, despite their ability to inhibit the uniculm phenotype in *cul2* mutants, single mutants with strong *eli-a* alleles are low tillering and typically produce about half as many tillers as non-mutants (Okagaki *et al.* 2018). In contrast, mutations in *INTERMEDIUM-C* (*INT-C*) and *MANY NODED DWARF* (*MND*) *4/6* result in high tillering phenotypes. *INT-C* is an ortholog of the branching inhibitor *TEOSINTE BRANCHED1* (*TB1*) in maize and encodes a TB1, CYCLOIDEA (CYC), PROLIFERATING CELL NUCLEAR ANTIGEN FACTOR1/2 (TCP) transcription factor. Loss-of-function *int-c* mutants have intermediate spike row-type (between 2-row and 6-row) and a moderate high tillering phenotype (Lundqvist and Lundqvist 1988; Ramsay *et al.* 2011). *MND 4/6* encodes a cytochrome P450 in the CYP78A family homologous to rice *PLASTOCHRON1* (*PLA1*), and *pla1* and *mnd* mutants both exhibit high rates of lateral organ initiation (Miyoshi *et al.* 2004; Mascher *et al.* 2014).

Quantitative trait loci (QTL) associated with tiller number have been found in coincident locations with genes regulating photoperiod sensitivity or spike row-type (Laurie *et al.* 1995; Karsai *et al.* 1997; Wang and Chee 2010; Naz *et al.* 2014; Alqudah *et al.* 2016; Nice *et al.* 2017). Photoperiod sensitivity in barley is largely determined by variation in *PHOTOPERIOD-H1*, an ortholog of *Arabidopsis PSEUDO RESPONSE REGULATOR 7* (*PRR7*). Plants with a dominant allele (*Ppd-H1*) are typically photoperiod sensitive and flower in response to long days, and plants with recessive alleles (*ppd-H1*) are typically photoperiod insensitive (Turner *et al.* 2005; Digel *et al.* 2015). Photoperiod sensitivity in barley is also influenced by variation in other genes, including *VERNALIZATION-H3* (*VRN-H3*) (Yan *et al.* 2006; Faure *et al.* 2007; Loscos *et al.* 2014), *VRN-H1* (Zitzewitz *et al.* 2005; Loscos *et al.* 2014), several *CONSTANS*-like genes (Campoli *et al.* 2012a; Mulki and von Korff 2016), and the barley ortholog of *Antirrhinum CENTRORADIALIS* (*HvCEN*) (Comadran *et al.* 2012). Photoperiod sensitivity impacts tiller number through influencing the timing and duration of shoot elongation, as tillering typically stops shortly after shoot elongation begins (García del Moral and García del Moral 1995; Miralles 2000). The influence of spike row-type on tiller number is usually attributed to a finite pool of resources that can be allocated to different developmental processes (Kirby and Jones 1977). Barley spikelets contain three florets, one central and two lateral, all of which are fertile and produce seeds in six-row barley (6-rows); whereas in two-row barley (2-rows) only the central floret is fertile. As a consequence of increased lateral spikelet fertility, 6-rows produce more, often smaller seeds than 2-rows, and they also tend to produce fewer tillers (Alqudah and Schnurbusch 2014, 2015; Liller *et al.* 2015). Spike row-type is primarily determined by variation in *SIX-ROWED SPIKE 1* (*VRS1*), which encodes a homeodomain leucine zipper protein (Komatsuda *et al.* 2007), or *VRS4*, which encodes an ortholog of the maize transcription factor RAMOSA2 (Koppolu *et al.* 2013), both of which are inhibitors of lateral spikelet development. Plants with dominant *VRS1* or *VRS4* alleles are typically 2-rows, whereas plants with recessive alleles are typically 6-rows. Variation in other genes that influence inflorescence morphology, including *VRS3* (van Esse *et al.* 2017; Bull *et al.* 2017) and *INTERMEDIUM* genes (Lundqvist and Lundqvist 1988; Ramsay *et al.* 2011), have also been shown to influence tiller number (Liller *et al.* 2015).

To date, most studies on the genetic control of tillering in barley have used forward genetics, as with the previously mentioned tillering mutants, or bi-parental mapping approaches (e.g. Arifuzzaman et al., 2014; Gyenis et al., 2007), which limit detection of natural genetic variation and the number of alleles that can be examined. However, a recent genome-wide association study identified QTL associated with tiller number at five developmental stages in a mapping panel of diverse spring barley accessions, and they showed genetic interactions between tiller number and spike row-type and photoperiod sensitivity (Alqudah *et al.* 2016). However, as this study was conducted in a greenhouse, the number of tillers that could be achieved, especially by high tillering accessions, was likely limited compared to field-grown barley.

In our study, a mapping panel consisting of 384 2-row and 384 6-row spring barley accessions from the National Small Grain Core Collection was examined. To increase tillering capacity, the panel was grown in the field and data on tiller number and rate and agronomic and morphological traits were obtained. To identify genetic variation associated with tiller number and developmental rate, the panel was genotyped using Genotyping-By-Sequencing (GBS) and a 50K SNP array (Bayer *et al.* 2017). Our objectives were to (1) quantify the genetic interactions between tillering and spike row type and photoperiod sensitivity; (2) identify potential trade-offs between tiller number and agronomic and yield-related traits; and (3) genetically map natural genetic variation associated with tillering and characterize the extent to which it overlaps genetic variation associated with related traits.

## MATERIALS AND METHODS

### Line Selection, Field Design, and Growing Conditions

A diversity panel containing 768 accessions (Table S1) from the National Small Grains Core Collection was developed for phenotypic analyses and genome-wide association studies (GWAS). The panel, split equally between 2-rows and 6-rows, was selected first by including the parents of a barley nested association mapping (NAM) population (Hemshrot et al., 2019; Smith, unpublished results) and then based on their contribution to polymorphism information content (PIC), as determined by Muñoz-Amatriaín et al. (2014). All accessions grown in 2014 and 2015 were the same except for seven lines that did not flower in 2014 were replaced with different lines in 2015.

The panel was grown in the field in St. Paul, MN in 2014 and 2015 in a Type 2 modified augmented design (Lin *et al.* 1983; Lin and Poushinsky 1985; May *et al.* 1989) containing 56 blocks, with one half containing 2-rows and the other half containing 6-rows (Figure S1). Individual blocks contained 15 rectangular 1.5 m by 0.3 m plots (five plots by three plots), with the central plot always containing a primary repeated check, cv. Conlon for 2-rows and cv. Rasmussen for 6-rows (Figure S1). Eight, randomly chosen blocks also contained two repeated secondary checks, assigned randomly to plots within the block. PI584962 and PI614939 were used as secondary checks for 2-rows, and PI327860 and CIho7153 were used as secondary checks for 6-rows. All other plots contained one of the 768 accessions from the mapping panel. To confirm trait correlations with tiller number and other traits from the 2014 and 2015 trials, in 2016, 54 lines split equally between 2-rows and 6-rows, were randomly chosen from NAM parent accessions grown in both years using the sample function in R (Table S1). The 54 accessions and the primary checks Conlon and Rasmussen were grown in a complete, randomized block design with three replicates. In all years, adjacent plots of non-vernalized winter wheat separated plots containing barley to control weeds, prevent shading, and allow space for lodging. Plots containing barley were machine planted with 30 seeds per plot and one week after emergence were thinned to ten plants per 1.5 m-long plot with regular spacing between plants.

### Phenotyping, trait value adjustment, and phenotypic analyses

Vegetative traits measured included tiller number, plant height, leaf width (2015 only), and stem diameter (2014 and 2015 only). In 2014 and 2015, tillers were counted on the same plants (ten in 2014 and five in 2015) per row weekly, beginning at two weeks past-emergence (2WPE) and ending at 7WPE. Productive tillers, tillers with grain-bearing spikes at plant maturity, were counted after grain filling when plants first showed signs of senescence (yellowing of awns and flag leaves). Tillering rate was calculated by dividing the maximum tiller number by the time in weeks that maximum tiller number occurred. Other metrics of tillering rate were determined by calculating the differences between mean tiller number between two consecutive weeks and by calculating the slope of a line fit to mean tiller number between at least three consecutive weeks. Leaf width (2015 only) and plant height were measured at the same time that productive tillers were counted. Plant height was calculated as the mean height (cm) of the tallest shoots of all plants from soil level to the top of the spike, not including the awns. Leaf width was calculated as the mean width (mm) at the widest point of the second leaf below the flag leaf on the tallest shoot of all plants. This leaf was chosen because it was consistently green at maturity. The tallest stem of all individual plants in a row were harvested after senescence and dried in an oven at 37 °C for 72 hours. Dried stems were scanned, and the diameters (mm) were measured at the widest point of the last internode (below the peduncle) and averaged for each accession using Image J software (version 1.50).

Inflorescence-related traits included spike row-type, seeds per spike, spike length, and 50-kernel weight. Spikes from the tallest shoots of five plants were harvested after senescence and dried in an oven at 37 °C for 72 hours. Spike length was measured from the base to the tip of the spike, not including awns. All seeds from the five spikes were removed by hand and counted; and mean seeds-per-spike was calculated. All seeds from the five spikes were pooled together, and 50-kernel weight was calculated as the total mass (g) divided by the total number of seeds multiplied by 50.

Days to heading was recorded when spikes on at least half of the shoots in a row were at least 50% emerged from the boot. Lodging was scored after senescence but before spikes were harvested, based on a scale of one to five, with one being completely upright and five being completely prostrate.

Trait values were adjusted using two different methods developed by Lin et al. (1983) specifically for Type 2 modified augmented designs and then assessed before and after correction to determine whether adjustment reduced heterogeneity of checks. One method, based on row and column averages of primary checks (Method 1 – M1), is better for correcting values when the field varies across plot rows and/or columns (Lin *et al.* 1983). Another method, based on linear regression of primary and secondary checks (Method 3 – M3), is better for correcting values when the field varies in many directions. M1 adjusted trait values (*M1AdjValue*) were calculated using the following equation:

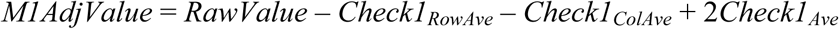

*Check1_RowAve_* and *Check1_ColAve_* were the averages of all primary check trait values in the same block row and block column, respectively, as the raw trait value being adjusted. *Check1_Ave_* was the average of all primary check values. Method 3 adjusted trait values (*M3AdjValue*) were calculated using the following equation:

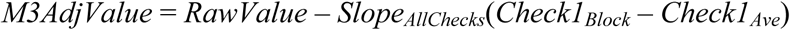

*Slope_AllChecks_* was the slope resulting from linear regression of primary check trait values versus the average secondary check trait values within the same block, and *Check1_Block_* is the value of the primary check in the same block as the raw trait value. Appropriateness of correction and selection of a correction method was based on two criteria (Lin and Poushinsky 1983, 1985; Lin et al., 1983; May et al., 1989). First, ANOVA in R (version 3.4.4) using primary check trait values was used to test for block row and column effects (Table S2). Second, relative efficiency of correction was calculated by dividing the average variance of raw secondary check trait values by the average variance of adjusted secondary check trait values, and values greater than one indicated that correction reduced variance due to heterogeneity in the field (Table S2). Raw trait values (Table S3) were used for phenotypic analyses to prevent individual trait adjustments from affecting trait correlations, and raw or adjusted (if applicable) trait values were used for genome-wide association mapping (Table S4).

All statistical analyses and data visualizations were performed in R. Broad-sense heritability (H^2^) was estimated using 2014 and 2015 raw trait values by two-way ANOVA with the following model: Trait ∼ Year + Line. Genetic variance was calculated as the difference between the line sum of squares and the residual sum of squares divided by two (for two years – 2014 and 2015), and heritability was calculated by dividing genetic variance by the sum of genetic variance and the residual sum of squares divided by two (Table 1 and Table S5). Estimates were based on lines that had trait data in both years, which varied depending on the trait, and the number of lines used for each trait estimate is included in Table 1 and Table S5. Trait heritability was also estimated with 2016 raw trait values using rep instead of year in the two-way ANOVA model (Table S5).

**Table 1.** Summary statistics for tillering traits measured in 2014 and 2015.

| Trait | Lines | Number of Lines | Mean |  | Standard Deviation |  | p-value<sup>a</sup> | H<sup>2</sup> |
| --- | --- | --- | --- | --- | --- | --- | --- | --- |
|  |  |  | 2014 | 2015 | 2014 | 2015 |  |  |
| Tiller Number 2WPE | All | 756 | 1.58 | 3.13 | 0.81 | 0.74 | 2.0E-09 | 0.35 |
|  | 2-Row | 375 | 1.84 | 3.38 | 0.83 | 0.76 | 0.032 | 0.17 |
|  | 6-Row | 381 | 1.31 | 2.88 | 0.70 | 0.62 | 3.7E-03 | 0.24 |
|  | *Ppd-H1* | 324 | 1.40 | 3.03 | 0.74 | 0.67 | 7.1E-05 | 0.35 |
|  | *ppd-H1* | 432 | 1.72 | 3.21 | 0.83 | 0.77 | 5.3E-05 | 0.31 |
| Tiller Number 3WPE | All | 756 | 3.39 | 8.00 | 1.63 | 2.26 | 6.4E-07 | 0.30 |
|  | 2-Row | 375 | 3.77 | 9.27 | 1.66 | 2.13 | 0.168 | 0.09 |
|  | 6-Row | 381 | 3.00 | 6.75 | 1.52 | 1.58 | 0.047 | 0.16 |
|  | *Ppd-H1* | 324 | 3.06 | 7.44 | 1.56 | 1.93 | 6.3E-04 | 0.30 |
|  | *ppd-H1* | 432 | 3.63 | 8.42 | 1.65 | 2.39 | 3.9E-03 | 0.23 |
| Tiller Number 4WPE | All | 756 | 5.66 | 12.98 | 2.33 | 3.58 | 6.2E-11 | 0.38 |
|  | 2-Row | 375 | 6.08 | 14.13 | 2.26 | 3.47 | 9.4E-04 | 0.28 |
|  | 6-Row | 381 | 5.24 | 11.85 | 2.34 | 3.33 | 2.7E-06 | 0.37 |
|  | *Ppd-H1* | 324 | 5.17 | 12.40 | 2.26 | 3.46 | 1.1E-07 | 0.44 |
|  | *ppd-H1* | 432 | 6.03 | 13.42 | 2.32 | 3.61 | 3.0E-04 | 0.28 |
| Tiller Number 5WPE | All | 756 | 7.11 | 19.27 | 2.98 | 6.28 | 3.0E-16 | 0.45 |
|  | 2-Row | 375 | 7.53 | 21.51 | 2.98 | 5.76 | 6.0E-06 | 0.37 |
|  | 6-Row | 381 | 6.68 | 17.06 | 2.92 | 5.98 | 1.8E-11 | 0.50 |
|  | *Ppd-H1* | 324 | 6.39 | 17.75 | 2.88 | 6.10 | 2.3E-11 | 0.52 |
|  | *ppd-H1* | 432 | 7.64 | 20.40 | 2.94 | 6.17 | 1.1E-05 | 0.34 |
| Tiller Number 6WPE | All | 756 | 7.21 | 19.97 | 3.28 | 6.73 | 4.4E-25 | 0.53 |
|  | 2-Row | 375 | 7.99 | 22.47 | 3.29 | 6.14 | 3.4E-08 | 0.43 |
|  | 6-Row | 381 | 6.45 | 17.51 | 3.10 | 6.38 | 1.3E-15 | 0.56 |
|  | *Ppd-H1* | 324 | 6.16 | 18.22 | 3.03 | 6.40 | 4.1E-15 | 0.58 |
|  | *ppd-H1* | 432 | 8.00 | 21.28 | 3.25 | 6.68 | 3.1E-09 | 0.43 |
| Tiller Number 7WPE | All | 756 | 6.72 | 18.98 | 3.31 | 6.54 | 2.1E-22 | 0.51 |
|  | 2-Row | 375 | 7.53 | 21.90 | 3.31 | 6.07 | 1.8E-06 | 0.38 |
|  | 6-Row | 381 | 5.92 | 16.11 | 3.11 | 5.67 | 3.7E-15 | 0.55 |
|  | *Ppd-H1* | 324 | 5.60 | 17.13 | 3.03 | 5.87 | 5.4E-12 | 0.54 |
|  | *ppd-H1* | 432 | 7.56 | 20.37 | 3.26 | 6.68 | 9.0E-09 | 0.42 |
| Tiller Number Maximum | All | 756 | 8.01 | 20.90 | 3.45 | 6.88 | 6.3E-22 | 0.50 |
|  | 2-Row | 375 | 8.69 | 23.44 | 3.42 | 6.22 | 1.2E-06 | 0.39 |
|  | 6-Row | 381 | 7.34 | 18.40 | 3.34 | 6.59 | 3.1E-15 | 0.56 |
|  | *Ppd-H1* | 324 | 6.95 | 19.29 | 3.24 | 6.71 | 4.2E-14 | 0.57 |
|  | *ppd-H1* | 432 | 8.81 | 22.11 | 3.38 | 6.76 | 1.5E-07 | 0.39 |
| Tiller Number Productive | All | 744 | 6.14 | 13.17 | 3.14 | 4.29 | 4.7E-13 | 0.41 |
|  | 2-Row | 372 | 6.96 | 15.43 | 3.14 | 4.13 | 4.3E-03 | 0.24 |
|  | 6-Row | 372 | 5.32 | 10.90 | 2.92 | 3.09 | 9.9E-05 | 0.32 |
|  | *Ppd-H1* | 314 | 4.92 | 11.98 | 2.47 | 3.38 | 6.8E-05 | 0.35 |
|  | *ppd-H1* | 430 | 7.04 | 14.04 | 3.27 | 4.66 | 3.4E-05 | 0.32 |

<sup>a</sup>p-values indicate significance of genetic variance based on 2-way ANOVA.

One-way ANOVA was performed followed by a Tukey-Kramer test for pairwise comparison of trait means between different year, spike row-type, and photoperiod sensitivity groups; and the multcompLetters function (multcompView, version 0.1-7) was used to assign letters designating whether groups were significantly different based on false discovery rate (FDR)-adjusted p-values from the Tukey-Kramer test. Pearson and Spearman rank correlations between traits were calculated using the rcorr function (Hmisc, version 4.1-1) (Table 2 and Table S6). A distance matrix was calculated based on average weekly (two to seven weeks past-emergence) and productive tiller number, and principal coordinates analysis (PCoA) of the distance matrix was performed using the cmdscale R function. The first and second principal coordinates based on tiller number were used as traits in association mapping (Table S4).

**Table 2.**
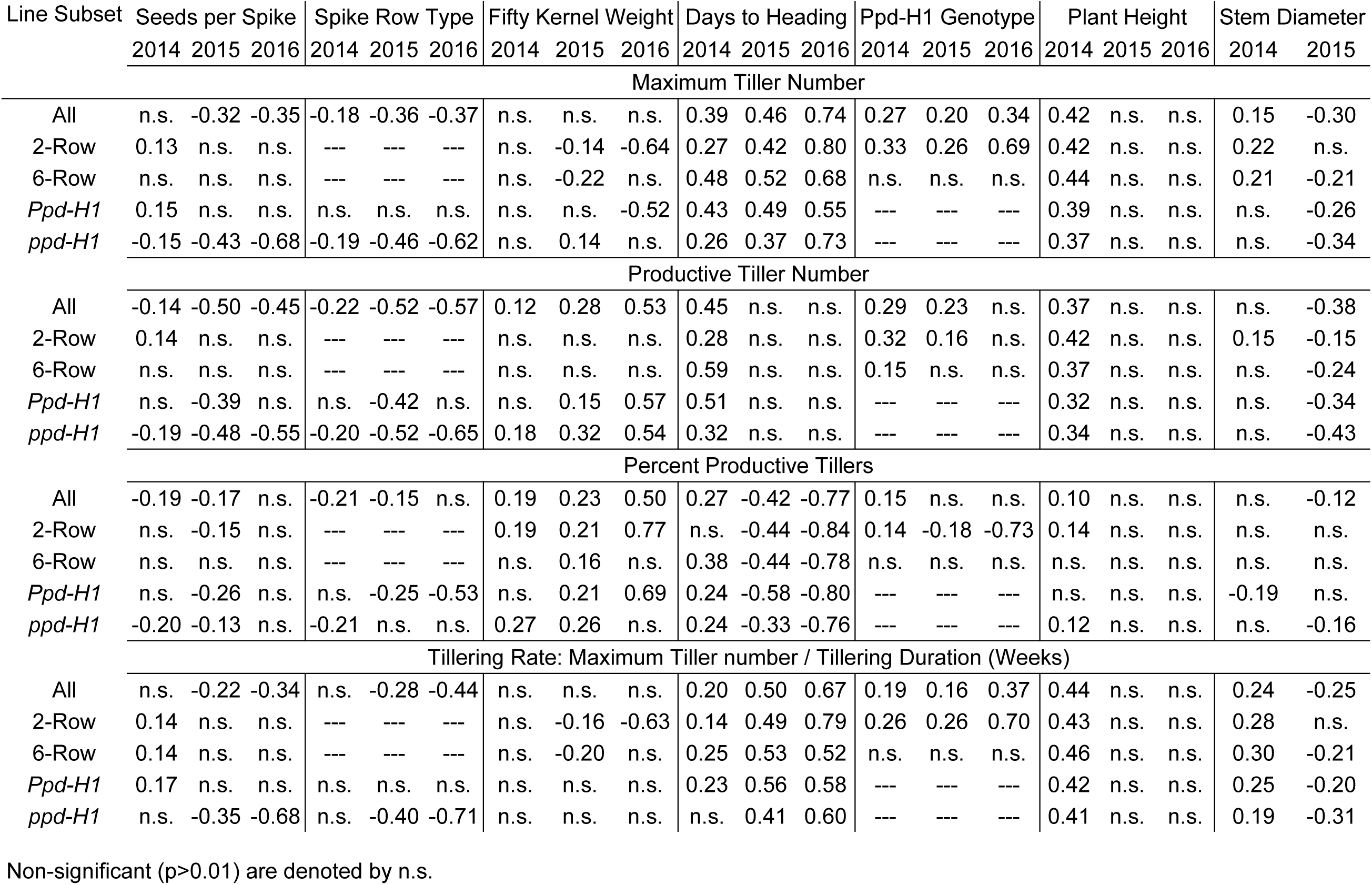
Pearson’s correlation coefficients for tillering traits versus other traits for lines grown in each year.

For multiple linear regression (MLR) analyses, the following model was fit using the lm function in R with tiller number as the response variable and other traits as predictor variables (File S1):

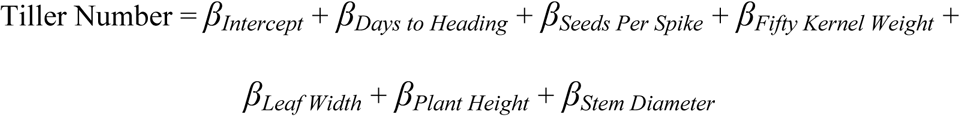

Before model fitting, lines with missing values for any of the traits included in the model were removed. The order of predictor variables in the MLR model was chosen based on relative contribution to R^2^, which was calculated using the “lmg” method (adapted Lindeman et al., 1980) from the boot.relimp function (relaimpo, version 3.3-2; Groemping, 2006). Next, the boot.stepAIC function (bootStepAIC, version 1.2-0) was used to choose a best-fit model by fitting the model 1000 times using forward and backward selection to choose predictor variables in the model. The final model was refit and outliers were removed based on Cook’s distance. Lines with the highest Cook’s distance were removed iteratively, and the model was refit until the R^2^ value of the model did not improve significantly. Predictor variables were checked for collinearity using the vif function (car, version 3.0-0) to ensure none of the variables had a Variance Inflation Factor (VIF) that indicated excessive correlation of predictor variables (VIF > 5). After all outlier lines were removed and the model was refit, the boot.relimp function was used to calculate relative proportion of total variance explained (contribution to R^2^ of the entire model) by individual predictor variables.

### Genotyping, Linkage Disequilibrium, and Population Structure Analysis

Lines were genotyped using GBS and a barley 50K iSelect SNP array (Bayer *et al.* 2017). DNA was extracted from seedling leaf tissue using a Mag-Bind® Plant DNA Plus kit (Omega Bio-tek, Norcross, GA), following the manufacturer’s instructions, and genomic DNA was quantified using a Quant-iT™ PicoGreen® dsDNA Assay Kit (Thermo Fisher Scientific, Waltham, MA). For GBS, reduced representation libraries were created according to Poland et al. (2012) using Pst1-Msp1 restriction enzymes. Libraries were sequenced using a HiSeq 2500 system (Illumina, San Diego, CA) to obtain single-end 125 bp reads. SNP calling was performed using the TASSEL 5 GBS Version 2 Pipeline using 64 base kmers and a minimum kmer count of five. Reads were aligned to the Morex reference genome assembly using the “aln” algorithm in the Burrows-Wheeler Aligner (BWA, version 0.7.10) (Mascher *et al.* 2017; Beier *et al.* 2017). Genotyping using barley 50K iSelect BeadChip kits (Illumina) was performed according to the manufacturer’s instructions, and SNPs were scored in GenomeStudio (version 2.0.2, Illumina) using manually curated clusters developed by Bayer et al. (2017). GBS and 50K SNP datasets were filtered individually based on percent missing data and percent heterozygosity. All filtering and imputing steps were performed using TASSEL 5. For the first round of filtering, GBS SNPs were removed if more than 50% of calls were missing or heterozygous and the minor allele frequency (MAF) was less than 0.03, and 50K array SNPs were eliminated if they contained more than 20% missing or heterozygous calls and a MAF less than 0.03. The GBS and 50K SNP datasets were then merged and missing data was imputed using the LD-kNNi imputation method in TASSEL 5 (sites = 20, Taxa = 5, maxLDDistance = −1). The merged, imputed SNP dataset was filtered again for missing data, eliminating SNPs and lines with more than 5% missing/heterozygous data. Lines were also filtered for missing data, and twenty-six lines with more than 5% missing/heterozygous SNP calls were excluded from association mapping and other genetic analyses. Three lines were removed from all genetic analyses because the spike row-type did not match what was recorded in GrainGenes (https://wheat.pw.usda.gov), GRIN (https://npgsweb.ars-grin.gov), and Muñoz-Amatriaín et al. (2014) (see notes in Table S1). SNPs were then tagged using the Tagger feature in Haploview (version 4.1) (Barrett *et al.* 2005) with an R^2^ cutoff of 0.95, resulting in 69,607 tagged SNPs for 747 lines (Table S7).

To analyze chromosomal linkage disequilibrium (LD) decay, pairwise R^2^ values between all SNPs within a chromosome were calculated using TASSEL 5, and the background LD level was calculated as the 95^th^ percentile of significant (pDiseq < 0.01) R^2^ values for all SNP pairs >= 50 cM apart, the distance at which the recombination rate is 0.5 for loci on the same chromosome. A non-linear model described by Hill and Weir (1988) was fit to all significant pairwise R^2^ values and their corresponding distances using the nls function in R, and the decay distance was calculated as the distance at which the non-linear model intersected with background LD level (Marroni *et al.* 2011) (File S2, S3). LD decay distances were calculated for individual chromosomes using physical and POPSEQ positions (Mascher *et al.* 2013; Beier *et al.* 2017) of tagged SNPs (Table S8). Based on LD decay distances, which were less than 1 cM for all chromosome (Table S8), a genetic distance of +/- 2 cM was chosen as a cutoff for including significant SNPs in the same quantitative trait loci (QTL) to account for regions with higher LD. To assess intrachromosomal patterns of LD for candidate gene analysis (as in Figure S8), pairwise comparisons were made between SNPs in 100 SNP windows. R^2^ values were ordered by mean position, and the R^2^ values and mean positions of 4950 pairwise comparisons (unique number of pairwise comparisons for 100 SNPs) were averaged and plotted as a line graph and a curve was fit using local regression (LOESS) (File S4).

Population structure was analyzed using the program STRUCTURE (version 2.3.4) (Pritchard *et al.* 2000). A set of 701 SNPs for STRUCTURE analysis (Table S9) were chosen by selecting SNPs from individual chromosomes from the final tagged SNP dataset that were at least as far apart as the calculated genetic decay distance (Table S8). Results from ten individual STRUCTURE runs for K 1-10 were analyzed using STRUCTURE Harvester (Earl and von Holdt 2012). The optimum number of subpopulations was chosen based on delta K (ΔK), which was calculated by STRUCTURE Harvester using equations from Evanno et al. (2005).

### Genome-wide Association Mapping

Genome-wide association mapping analysis was performed using compressed mixed linear models from the GAPIT R package (Genome Association and Prediction Integrated Tool, version 2.0) (Lipka *et al.* 2012) with the final imputed and filtered set of 69,607 SNP tags (Table S7) and raw and corrected (if applicable based on Table S2) phenotypic data (Table S4). The MAF cutoff was 0.03 for all lines (n=727-740, depending on the trait) and 0.05 for subsets based on spike row-type or *PPD-H1* alleles (n=305-437, depending on the subset and trait). The model selection feature of GAPIT was used to choose the optimum number of principal components for each individual trait to account for population structure, and the optimal compression level determined by GAPIT was used. The percentage of genetic variance explained by individual SNPs was calculated as the difference between R^2^ of models with the SNP and without the SNP. Information about all significant SNPs, including allelic effect size, percent variance explained, and nearest gene information is included in Table S10.

### Data Availability

All data necessary for reproducing results are available within supplemental tables, which are available in FigShare. Table S1 contains information about all accessions, including collection site, improvement status, spike row-type, and STRUCTURE subpopulation assignment. Table S7 contains all SNP markers used for association mapping, and Table S9 contains all SNP markers used for STRUCTURE analysis. Raw trait data used for phenotypic analyses is included in Table S3, and trait data used for association mapping is included in Table S4. Supplemental figures and R scripts for multiple linear regression and LD analyses (Files S1-S4) are also available in FigShare.

## RESULTS AND DISCUSSION

### Tiller number in the two- and six-row diversity panel

In 2014 and 2015, 761 lines were grown in the field, and data were collected for weekly and productive tiller number, days to heading, plant height, stem diameter, leaf width (2015 only), seeds per spike, fifty kernel weight, and lodging (2015 only) (Table S3). Fifty-four lines that were grown in 2014 and 2015 were also grown in 2016 in three complete, randomized blocks, and data for weekly and productive tiller number, days to heading, plant height, seeds per spike, and fifty kernel weight were collected (Table S3). Phenotypic data were analyzed in all lines and in subsets of lines based on spike row-type and *PPD-H1* alleles. Tiller number data from 2014 and 2015 are summarized in Table 1, and all trait data from all years are summarized in Table S5.

Genetic variance for tiller number was significant (p-value < 0.0001) in 2014, 2015, and 2016 for most time points (Table 1, Table S5). In both years for all line subsets, variance was highest for maximum tiller number and tiller number measured at later time points (5-7WPE), and it decreased for productive tiller number (Table 1). Tiller number at 6WPE, the time point at which maximum tiller number occurred on average for all lines, also had the highest heritability estimate (0.53) of all tiller counts. Decreased heritability from 6WPE to productive tiller number was likely due to variability in tiller survival, which appears to be strongly influenced by environment as genetic variance for percent productive tillers was not significant (Table S5). Heritability estimates for tillering traits were lower than other traits measured (Table S5).

Tiller number was compared using data for the 54 lines (27 2-rows and 27 6-rows) grown in all three years. Due to waterlogging in the field early in development in 2014, the onset of tiller development was delayed and maximum and productive tiller number was much lower than 2015 and 2016 (Figure 1A,B). By 2WPE in 2014, 25.4% of all lines grown had not yet developed at least one tiller per plant on average, whereas all lines grown in 2015 had developed at least one tiller per plant by 2WPE. Maximum tiller number was not significantly different between 2015 and 2016, but productive tiller number was lower in 2016 than 2015 due to lower tiller survival (Figure 1B). Despite differences between years, they all followed a similar trend where average tiller number increased linearly until 5WPE, after which it either slowed or began decreasing (Figure 1A).

**Figure 1.**
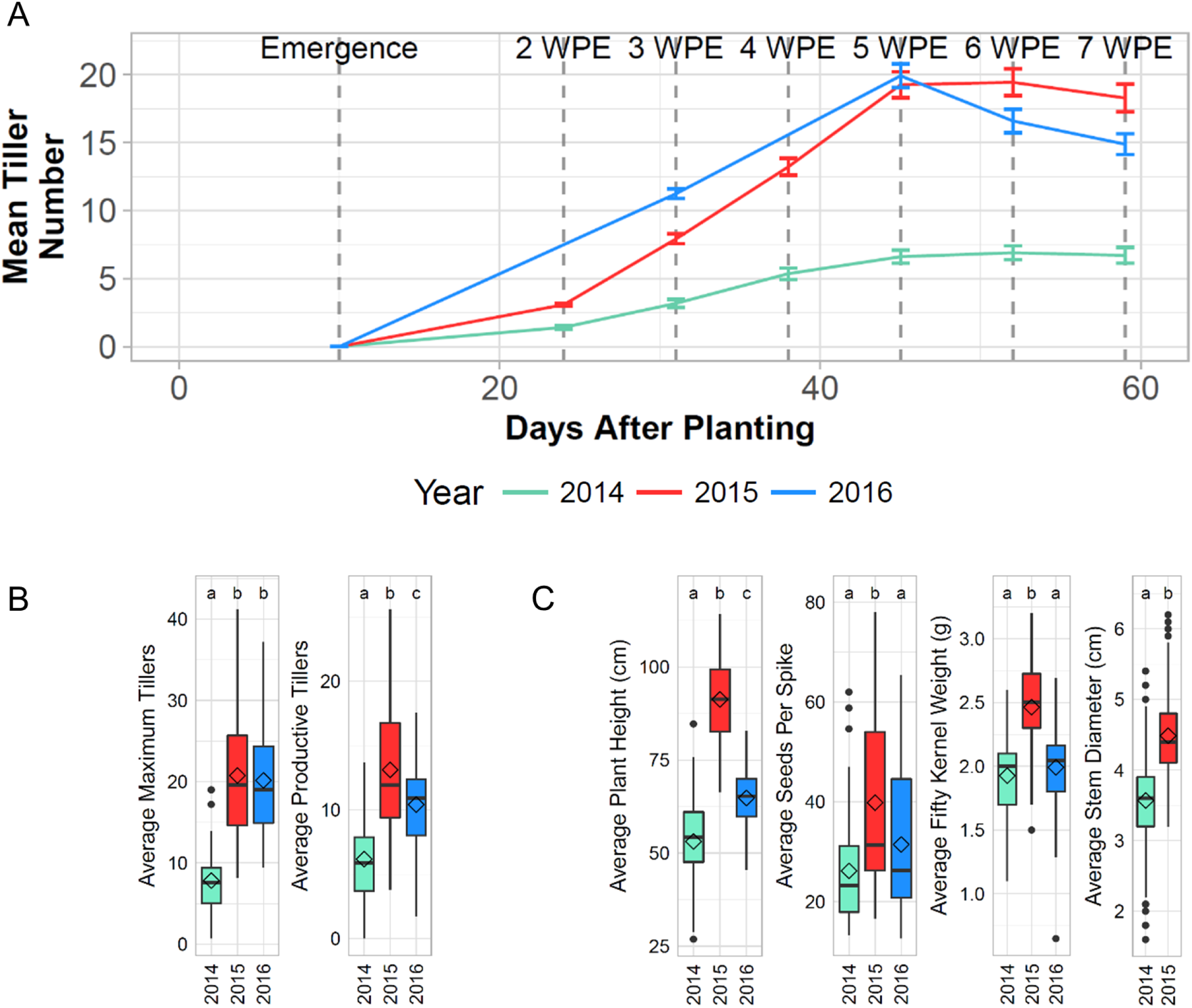
Overview of tiller development (tillering) and other traits. A) Progression of average tiller number throughout the growing season for 54 lines grown in 2014, 2015, and 2016. B) Box plots summarizing tillering traits for 54 lines grown in all three years. Diamonds represent mean trait values, and letters indicate whether groups are significantly different based on FDR-adjusted p-values from ANOVA in conjunction with Tukey Test. C) Box plots summarizing non-tillering traits show similar relationship between years as average productive tiller number.

Average plant height, stem diameter (measured in 2014 and 2015 only), seeds-per-spike, and fifty kernel weight followed a similar trend as productive tiller number across the three years, where trait values were highest in 2015 and lowest in 2014 (Figure 1C). In years when plants developed more productive tillers on average, they were also taller with thicker stems, more seeds per spike, and heavier seeds on average (Figure 1C), indicating that productive tiller number is correlated with overall plant fitness.

### Days to heading and spike row type explain a large proportion of variance in tiller number

Consistent with previous studies (Liller *et al.* 2015; Alqudah *et al.* 2016), our results support the observations that spike row-type and photoperiod response influence tiller number. However, these previous studies have not attempted to quantify the extent that these traits influence tiller number, nor have they assessed the simultaneous effects of both traits on tiller number. To gain a better understanding of these relationships, we examined tiller number in 761 lines in relation to days to heading, *PPD-H1* genotype, and spike row-type.

Spike row-type has been shown to influence tiller number as well as other traits like seed number and weight, and leaf area (Alqudah and Schnurbusch 2014, 2015; Liller *et al.* 2015). As expected, average tiller number was higher in 2-rows than 6-rows in 2014 and 2015 (Table 1). Duration of tiller development was also slightly longer for 2-rows than 6-rows in both years, and a lower percentage of tillers were productive in 6-rows compared to 2-rows in both years (Figure S2A). As commonly observed, most 2-rows also had thinner stems, narrower leaves, and longer spikes with fewer, heavier seeds than 6-rows (Figure S2B). Despite the difference in average tiller number, productive tiller number distributions in 2-rows and 6-rows largely overlapped (Figure S2C). Furthermore, some 6-rows produced as many tillers as high tillering 2-rows, and some 2-rows produced as few tillers as low tillering 6-rows (Figure S2C).

In earlier studies, variation in *PPD-H1* was shown to influence days to heading, leaf size, tiller number, and tillering duration (Turner *et al.* 2005; Alqudah *et al.* 2016, 2018; Digel *et al.* 2016). One SNP included in this study, BK_14, is 308 bp upstream of *PPD-H1* and has been previously shown to be in complete or near-complete LD with a SNP in the CONSTANS (CO), CO-like, and TOC1 (CCT) domain of *Ppd-H1* and is a likely causal variant underlying photoperiod sensitivity differences (Turner *et al.* 2005; Digel *et al.* 2016). LD analysis indicated that all SNPs in *PPD-H1* and several that flanked it were in high LD (Figure S3). Therefore, BK_14 was used to distinguish lines as having the photoperiod sensitive *Ppd-H1* (G) allele or the photoperiod insensitive *ppd-H1* (A) allele, and correlation of *PPD-H1* alleles and tiller number was assessed separately in 2-rows and 6-rows. We found that 2-row accessions carrying *ppd-H1* had more tillers than 2-rows carrying *Ppd-H1*, but tiller number was not significantly different between 6-rows carrying the two *PPD-H1* alleles (Figure S4A). Interestingly, days to heading explained a larger proportion of variance in multiple linear regression (MLR) models of tiller number in 6-rows than 2-rows in both years (Figure S4B), suggesting that variation in other genes that influence photoperiod sensitivity could affect tiller number more strongly than *PPD-H1* in this 6-row germplasm.

The large number of lines included in this study allowed us to characterize and quantify percent variance in tiller number explained by both spike row-type and photoperiod sensitivity simultaneously. Only data from 2015 was used for these analyses because more traits were measured in 2015 and variance in tiller number was higher than in 2014, as shown by higher standard deviation in tiller number (Table 1). In addition, photoperiod response was represented by days to heading in these analyses, and spike row-type was represented by seeds per spike in MLR models for all lines.

MLR models with tiller number as the response variable and other traits as predictor variables indicated that days to heading and spike row type explained a high proportion of variance in tiller number (Figure 2A). Together they explained 28% of the total variance in maximum tiller number and 12% of the total variance in productive tiller number (Figure 2A). Interestingly, a very small proportion of variance in productive tiller number was explained by days to heading (1.9%) (Figure 2A), probably due to variability in tiller survival between lines. Average differences in tiller survival represented by percent productive tillers between 2-rows and 6-rows (Figure S2A) could explain why seeds per spike accounted for a larger proportion of variance in productive tiller number than maximum tiller number.

**Figure 2.**
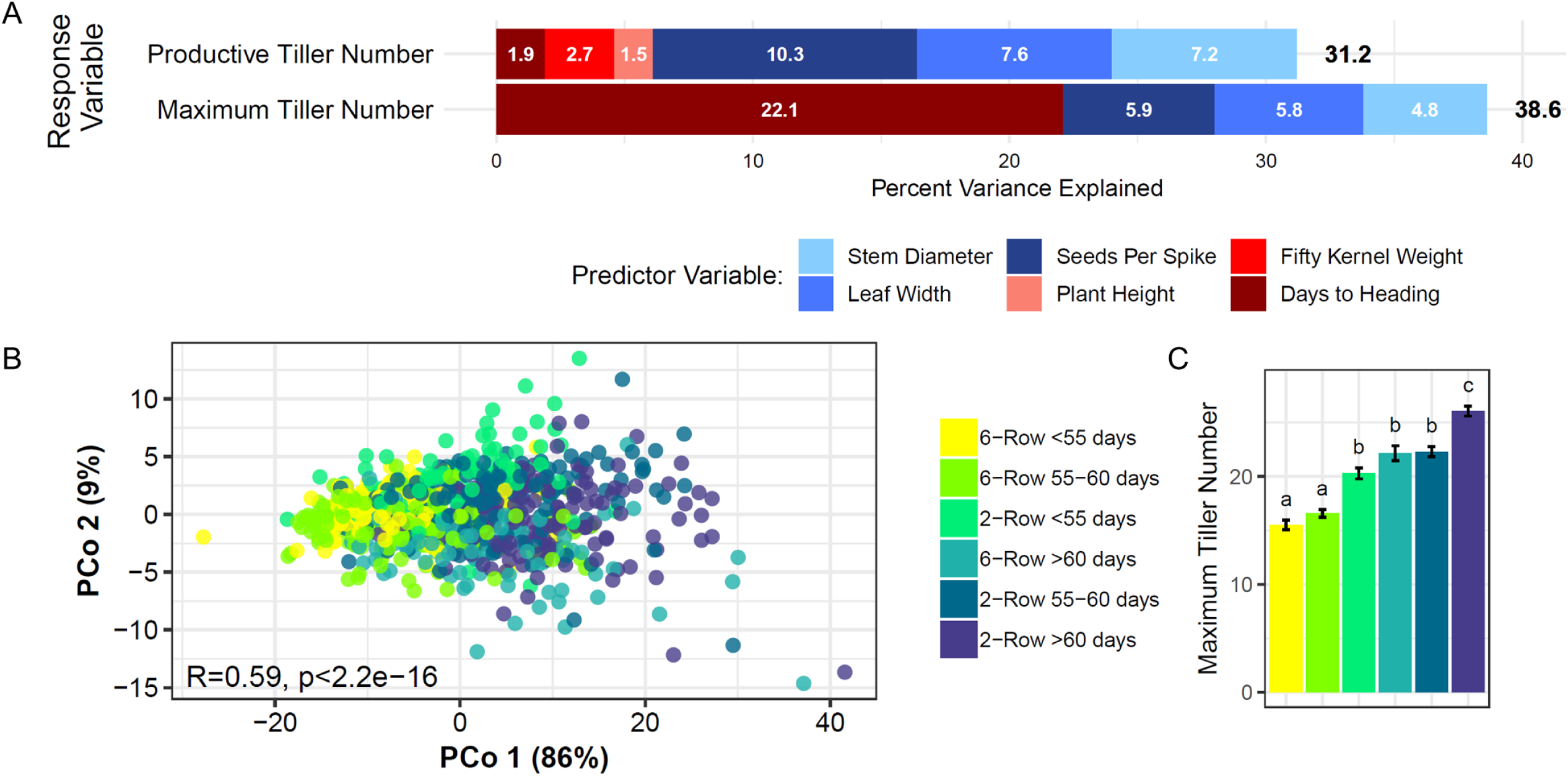
Days to heading and spike row-type explained a large proportion of variance in tiller number in 2015. A) Bar plots showing percent variance explained by predictor variables in multiple linear regression models of maximum and productive tiller number for all lines. White numbers on bars represent percent variance explained by individual predictor variables. Traits shaded in red and blue are positively and negatively associated with tiller number, respectively. Numbers beside bars are total percent variance explained (R^2^) of the entire model. Seeds per spike was included in the model as a proxy for spike row-type (2-row or 6-row). B) Principal coordinates (PCo) analysis based on weekly and productive tiller number measurements for all lines. Percent variance explained by PCo 1 and 2 is shown on axes. Strong correlation between PCo 1 and groups based on spike row-type (2-row or 6-row) and days to heading (R=0.59, p < 2.2e-16) indicates that these traits explain a large proportion of variance in tiller number. C) Comparison of mean tiller number at six weeks past emergence (WPE) between groups based on spike row-type and days to heading.

Principal coordinates (PCo) analysis based on tiller number throughout development and productive tiller number also indicated that a large proportion of variance in tiller number was explained by days to heading and spike row-type. Groups based on spike row-type and days to heading were more strongly correlated than any other single trait with PCo1 (R=0.59, p< 2.2e-16), which explained 86% of the total variance in the PCo model (Figure 2B). Furthermore, although 6-rows produced fewer tillers on average than 2-rows, maximum tiller number in late heading 6-rows (>60 days) was not significantly different from earlier heading 2-rows (<60 days), indicating that high tiller number can be achieved in late heading 6-rows (Figure 2C).

### Trade-offs between tillering and other traits

Tiller number and other traits were compared to evaluate trade-offs associated with high tiller number. Because spike row-type influences tiller number and other traits, trade-offs were assessed separately in 2-row and 6-row subsets and using 2015 data only for the same reasons as previously described. Results of MLR modeling indicated minor trade-offs between tiller number and other vegetative traits. Leaf width and stem diameter explained a significant proportion of variance in productive and maximum tiller number MLR models (Figure 3A), and their coefficients were consistently negative, indicating a tendency for leaf width and stem diameter to decrease as tiller number increased. Both traits were also weakly, negatively correlated with productive tiller number (Table 2 and Table S6).

**Figure 3.**
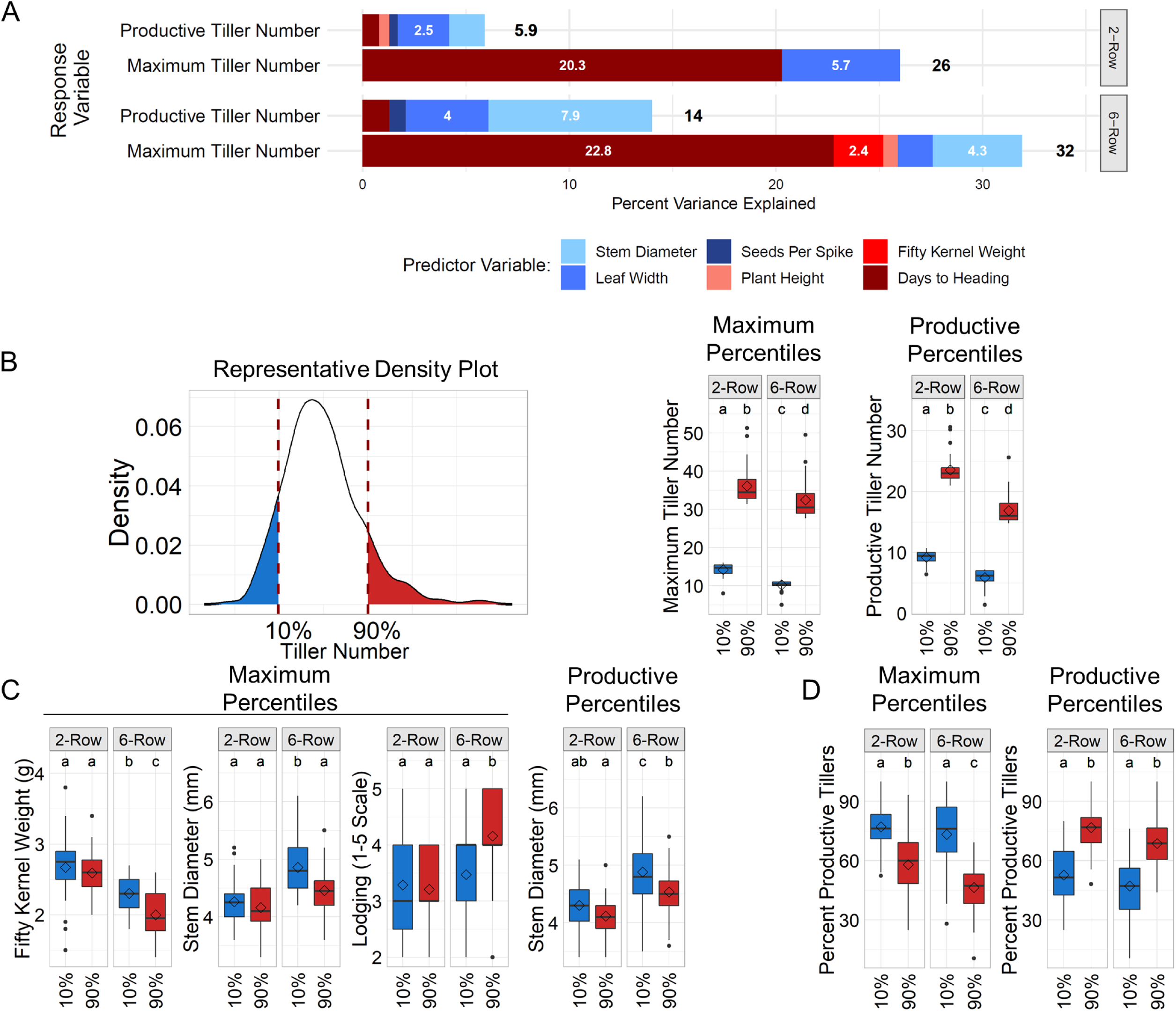
Minor trade-offs between tiller number and other traits in 2015. A) Percent variance explained by predictor variables in multiple linear regression models of maximum tiller number in 2-rows and 6-rows in 2015. White numbers on bars represent percent variance explained by individual predictor variables (< 2% if number is not shown). Traits shaded in red and blue are positively and negatively associated with tiller number, respectively. Numbers beside bars are total percent variance explained (R^2^) of the entire model. B) (Left) Representative density plot on left illustrates assignment of 2-row and 6-row lines into percentile groups based on maximum and productive tiller number. (Right) Comparison of tiller number in percentile groups based on maximum and productive tiller number. Diamonds represent mean trait values, and letters indicate whether groups were significantly different (Tukey Test, FDR-adjusted p-value < 0.01). C) Box plots showing traits that were significantly different between percentile groups based on maximum and productive tiller number. D) Box plots of percent productive tillers (tillers that survive and form grain-bearing spikes).

We considered the possibility that larger trade-offs or trade-offs that were not indicated by correlations or MLR modeling could be identified by comparing traits in lines with very different tillering capacities. Therefore, 2-rows and 6-rows were split into 10^th^ and 90^th^ percentile groups based on maximum and productive tiller number (Figure 3B). Despite at least 2.5-fold or higher change in average tiller number between percentile groups (Figure 3B), very few traits were significantly different between percentile groups. Stem diameter was lower in high tillering 6-rows (90^th^ percentile, maximum and productive) than low tillering 6-rows (10^th^ percentile, maximum and productive) but was not significantly different between high and low tillering 2-rows (Figure 3C). Fifty kernel weight was also lower and lodging severity increased in high tillering 6-rows (90^th^ percentile, maximum) than low tillering 6-rows (10^th^ percentile, maximum), but they were not significantly different between high and low tillering 2-rows (Figure 3C). Interestingly, the trend in percent productive tillers between percentiles based on maximum tiller number was reversed in percentiles based on productive tiller number (Figure 3D). This suggests that tiller survival had a major impact on final productive tiller number in 2015 and that variation in tiller survival may alleviate trade-offs between tiller number and other traits. Overall, our results suggest that trade-offs between tiller number and other traits were very minor and were slightly more pronounced in 6-rows than 2-rows, but, in general, there were no major trade-offs between tiller number and other traits independent of spike row-type.

It is likely that lower tiller number in 6-rows than 2-rows is due to a trade-off with seeds per spike, which is inherently higher in 6-rows (Figure S2B). However, there was no evidence from our study that more seeds per spike within 2-row or 6-row groups was associated with lower tiller number. Overall, results from this study indicated that trade-offs between tiller number and seeds per spike probably only exist if the difference in seeds per spike is very large, as it is between 2-rows and 6-rows.

Few studies have described trade-offs between tiller number and other traits in barley or other small grain crops, and the results have been inconsistent. For example, Kebrom et al. (2012) reported that removing tillers in wheat could induce development of larger spikes with more seeds. However, another study examined yield and yield-related traits in barley under different seeding densities over two years and found that there was no trade-off between tillers per plant and seeds per spike (Stoskopf and Reinbergs 1966). They found that the seeding density at which seeds per spike was highest was the same density at which productive tiller number per plant was highest. Furthermore, when they compared 20 high-yielding lines and 20 low-yielding lines, they found that average seeds per spike was higher in high-yielding lines but that average tiller number was not different.

### Natural genetic variation associated with tillering

Population structure was characterized in all lines in the diversity panel prior to association mapping. As with the entire NSGC collection, population structure analysis of all lines in the diversity panel using STRUCTURE resulted in five subpopulations, corresponding to those described in Muñoz et al. (2014), that were distinguished primarily by spike row-type, collection location, and improvement status (Figure S5 and Table S1). Days to heading and tiller number did not vary by improvement status (landraces versus cultivars) in Subpopulations (SP) 1, 3, and 4 (Figure S6). SP2 and SP5 were not compared because they almost exclusively contained landraces (Figure S5). Tiller number was higher in SP3 than SP1 or SP4 (Figure S6), but this was likely due to the fact that SP3 contained primarily 2-rows while SP1 and SP4 contained primarily 6-rows.

Genome-wide association mapping was performed using 2014 and 2015 raw or adjusted (if applicable based on Table S2) phenotypic data for all tillering traits, days to heading, and spike row-type. Tillering QTL included SNPs significantly associated with tiller number, rate of tillering, and tillering principal coordinates. Tiller number included 2-7WPE, productive, and maximum tiller number. Thirty-seven QTL were associated with tillering traits in 2014 and 2015, (Table 3); however, only four were identified in both years, one on 2H at 56.82-58.76 cM (2H-58), one on 5H at 47.89-48.10 cM (5H-48), and two on 7H at 31-33.67 cM (7H-33) and 70.16-70.54 cM (7H-70) (Table 3, Figure 4A). These four tillering QTL accounted for a very small proportion of variance in tillering traits (Table S10), while the QTL that explained the most variance in tiller number were not detected in both years, one on 2H at 13.72 – 23.24 cM (2H-19) in 2014, and one on 3H at 35.39 cM (3H-135) in 2015 (Figure S7). The 2H-19 QTL overlapped the *PPD-H1* locus and was associated with tiller number, tillering rate, and tillering PCo1 in all lines and with tiller number and tillering rate in 2-rows (Figure S7). For many tillering traits in 2014, 2H-19 was the only QTL identified (Figure S7), and the allelic effect size for tiller number measurements ranged from 1.1-1.5 tillers (Table S10). The 3H-135 QTL was associated with tiller number, tillering rate, and tillering PCo1 in all lines and *Ppd-H1* lines, and with tiller number and tillering rate in 6-rows (Figure S7). For many tillering traits in 2015, 3H-135 was the only QTL identified, and the allelic effect size for tiller number measurements ranged from 1.5-4 tillers (Table S10).

**Figure 4.**
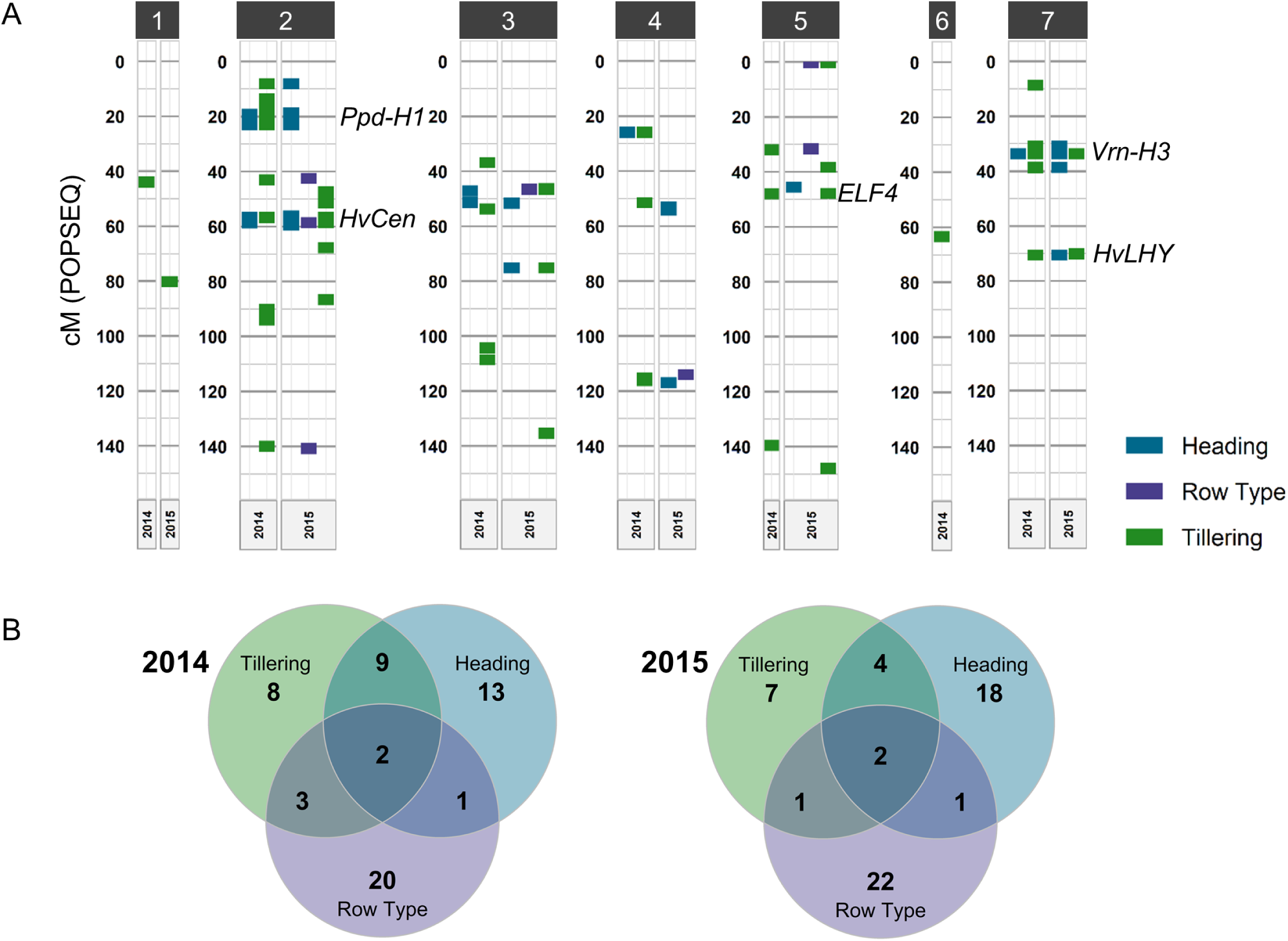
Most quantitative trait loci (QTL) associated with tillering overlapped QTL associated with days to heading and/or spike row-type. (A) Genetic positions on all chromosomes of significant SNPs (+/- 2 cM) associated with tillering, days to heading, and spike row-type. Only heading and row-type QTL that overlapped tillering QTL are shown. (B) Venn diagrams showing the number of tillering QTL in 2014 and 2015 that overlapped QTL associated with days to heading and spike row-type.

**Table 3.**
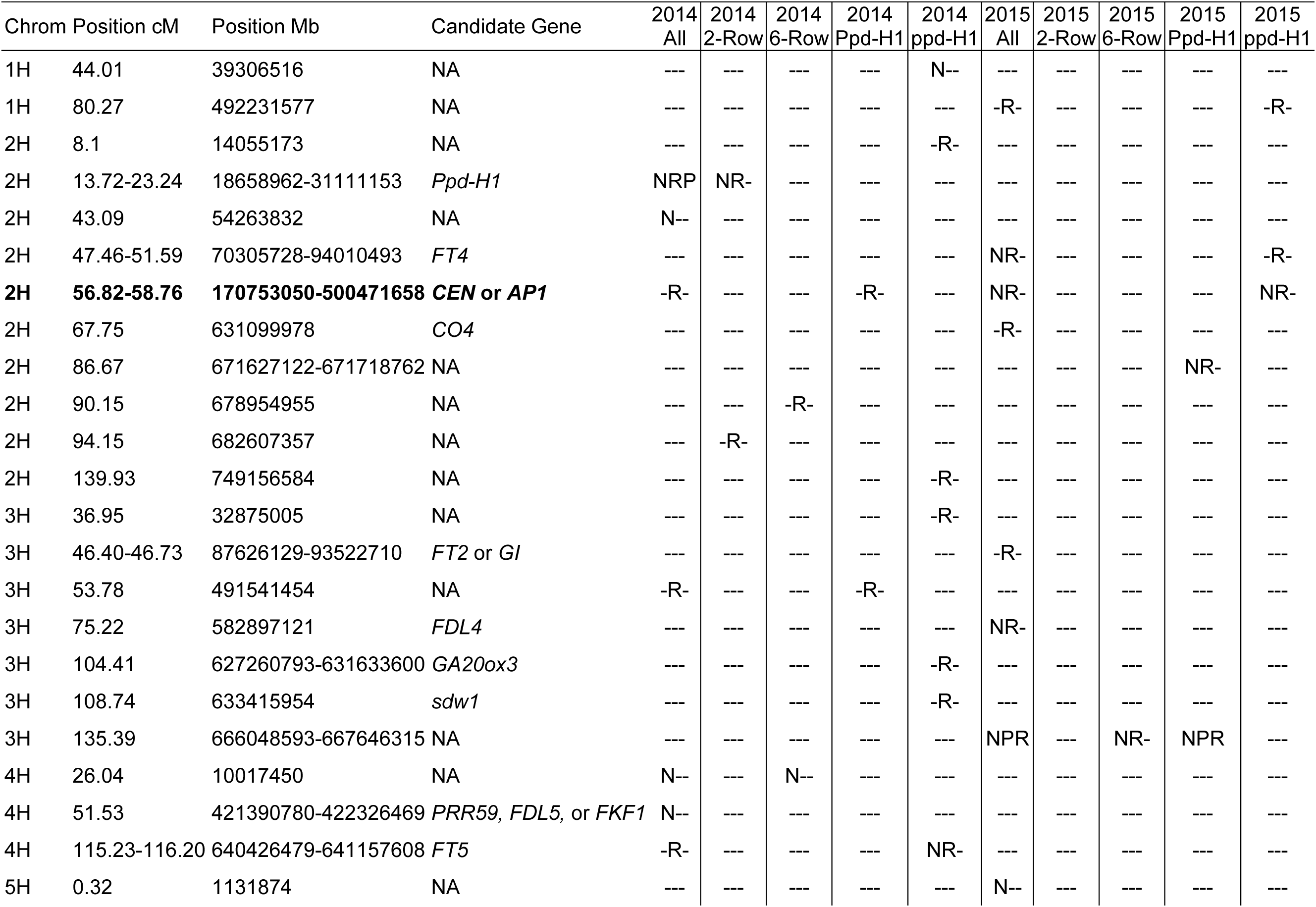
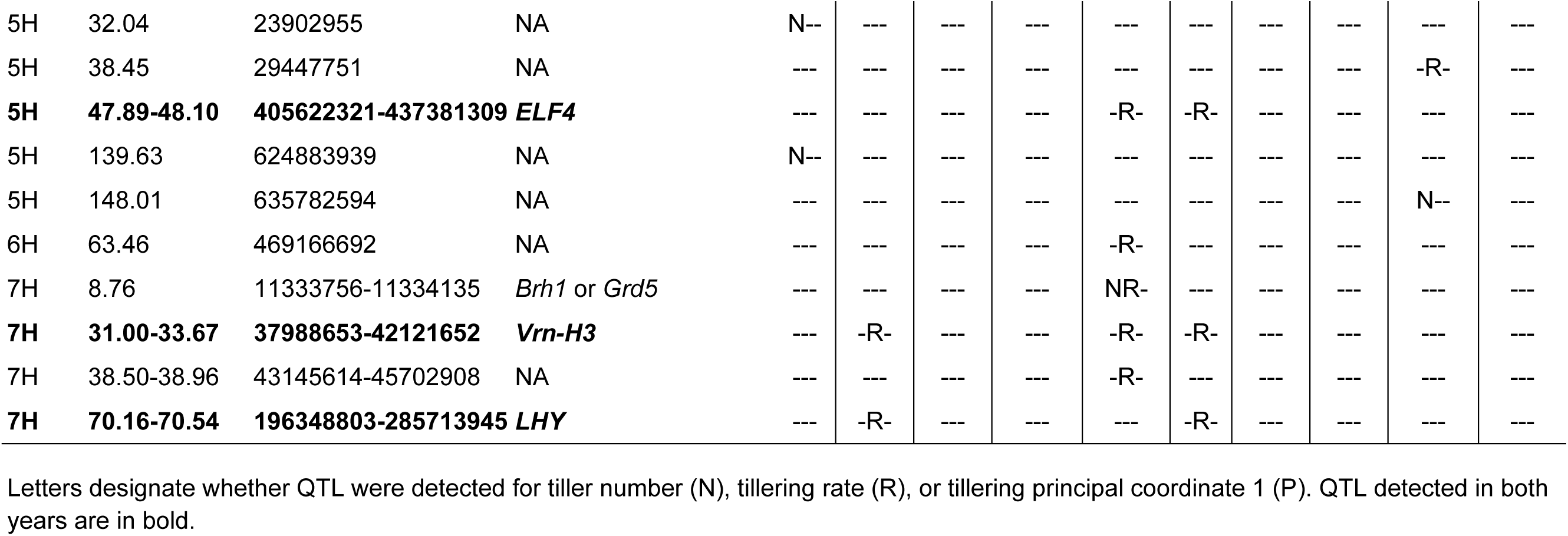
Quantitative trait loci (QTL) associated with tillering in 2014 and 2015.

Measuring tiller number throughout development provided opportunities to identify QTL associated with tillering rate, and to compare the number of QTL associated with tillering at different time points. Fourteen out of 23 and six out of 14 tillering QTL were associated only with tillering rate, and not tiller number, in 2014 and 2015, respectively (Table 3). Tiller number at later time points (5-7WPE, maximum, and productive) was associated with more QTL than at earlier time points (Figure S7). No QTL were associated with tiller number at 2WPE in either year; and no QTL were associated with tillering rate early in development (2-4 WPE) in 2014 (Figure S7), possibly due to low phenotypic variance during seedling development (Table 2).

Grouping lines based on their *PPD-H1* genotype and spike row-type allowed us to identify QTL that were not identified in all lines, and to observe that there was virtually no overlap in QTL detected in 2-rows and 6-rows or *Ppd-H1* and *ppd-H1* lines. In 2014, very few (four out of the 23) QTL associated with tillering were uniquely identified in all lines, whereas ten unique tillering QTL were identified in *ppd-H1* lines (Figure S7). Two QTL were uniquely identified in 2-rows and one was uniquely identified in 6-rows in 2014 (Table 3). No unique QTL were identified in *Ppd-H1* lines in 2014 (Table 3). In 2015, more QTL were identified in all lines than in any other group. All of the tillering QTL identified in 6-rows were also identified in all lines, and despite high phenotypic variance in 2015, no QTL were associated with tillering in 2-rows, possibly due to low allele frequency and the presence of many small effect loci that influence tillering. Including the *Ppd-H1* group enabled identification of three unique QTL (Table 3). In addition to identifying unique QTL within each year, including groups based on spike row-type and *Ppd-H1* genotype also enabled detection of three of the four QTL that were associated with tillering in both years. Only one of the four tillering QTL identified in both years, 2H-58, was identified in all lines in both years (Table 3).

Interestingly, three of the four QTL identified in both years in this study were also identified in a study by Alqudah et al. (2016) (2H-58, 5H-48, and 7H-70), which measured tiller number throughout development in a greenhouse-grown diversity panel, suggesting that these three QTL consistently influence tiller number under different environmental conditions. In total, ten of the 33 tillering QTL identified in this study were also identified in the Alqudah et al. study – relatively few considering the large number of QTL identified between the two studies. This modest overlap could be attributed to differences in overall tillering capacity between greenhouse-grown and field-grown barley, as field-grown barley has more potential to reach higher tillering capacities under favorable conditions. This could also explain the low overlap between the two years in our study, as tillering capacities differed greatly between the two years. It is also possible that the different diversity panels used in our study and the Alqudah et al. study harbor different alleles that influence tiller number. Therefore, growing different mapping panels under different environmental conditions is probably necessary to capture the full extent of natural genetic variation underlying tiller development.

### Overlap of natural genetic variation associated with tillering, days to heading, and spike row-type

Because tiller number was correlated with days to heading and spike row-type, we expected to see some overlap between QTL associated with these traits. In 2014, nine of 23 tillering QTL were also associated with row-type and/or heading, and in 2015, seven out of 14 tillering QTL were also associated with row-type and/or heading (Figure 4A). However, if all QTL associated with heading regardless of year were included, overlap between tillering QTL and heading QTL, especially in 2014, was much more extensive (Figure 4B). Incidentally, there was very little overlap between row-type QTL and heading QTL in either year (Figure 4B). Only one tillering QTL, 2H-58, which was the only one associated with tillering in all lines in both years, was also the only one associated with heading and row-type in both years (Figure 4A).

Interestingly, all four of the tillering QTL identified in 2014 and 2015 overlapped genes that have been previously shown to influence heading or circadian rhythm in barley, and all of them were also associated with heading in this study (Figure 4A). *HvCEN* (HORVU2Hr1G072750, 58.7 cM) is located in the 2H-58 QTL interval (Table 3) and was shown in a recent study that characterized 23 independent *HvCEN* mutants to influence flowering time, the number of spikelets per spike, and tiller number (Bi *et al.* 2019). Variation in *HvCEN* was also associated with days to heading in earlier studies (Comadran *et al.* 2012; Loscos *et al.* 2014). As previously mentioned, QTL in this region were identified for tiller number, days to heading, and spike row-type in all lines in both years. Although variation in *HvCEN* affects the number of spikelets per spike, there is no evidence that it affects the number of fertile florets per spikelet, so it is likely that another gene in this region is associated with spike row-type. *HvMADS15*, a MADS-box gene homologous to *APETALA1/FRUITFULL* (HORVU2Hr1G063800, 58.76 cM) is a more likely candidate because its expression is nearly undetectable in spike row-type *vrs3/int-c* double mutants, indicating a role in spike row-type determination (Zwirek *et al.* 2019). *VRS3* encodes a histone demethylase, and mutants have an intermediate spike row-type like *int-c* mutants (van Esse *et al.* 2017; Bull *et al.* 2017). The 5H-48 QTL overlaps *HvELF4-like* (HORVU5Hr1G060000, 48.4 cM), a homolog of Arabidopsis *EARLY FLOWERING 4* that is a likely candidate for environmental adaptation selection in barley landraces (Russell *et al.* 2016). *HvFT1*/*VRN-H3* (HORVU7Hr1G024610, 33.67 cM), an ortholog of Arabidopsis *FLOWERING LOCUS T* (*FT*), is located in the 7H-33 QTL interval and is an important regulator of flowering time in barley. Russell et al. (2016) found that *HvFT1* was more strongly associated with latitude in landraces than any other flowering gene, indicating its importance for adaptation, and variation in *HvFT1* was associated environmental adaptation and days to heading in other studies as well (Casas et al., 2011; Loscos et al., 2014; Maurer et al., 2015). The fourth QTL identified in both years for tillering rate and heading, 7H-70, co-localized with a probable ortholog (HORVU7Hr1G070870, 70.8 cM), based on sequence homology and circadian expression pattern, of the partially redundant circadian genes in Arabidopsis, *CIRCADIAN CLOCK ASSOCIATED 1* (*CCA1*) and *LATE ELONGATED HYPOCOTYL* (*LHY*) (Campoli *et al.* 2012b).

We found that more tillering QTL colocalized with days to heading QTL than with spike row-type QTL (Figure 4B), and surprisingly, no tillering QTL overlapped the *VRS1* locus or other *VRS* loci in either year, despite significant differences in all tillering traits between 2-rows and 6-rows in both years. This could be due to the extensive overlap in tiller number distributions between 2-rows and 6-rows that was previously mentioned (Figure S2C).

### Tillering QTL do not overlap known tillering genes

As previously described, mutations influencing tiller number have been identified and several mutated genes have been characterized. Interestingly, none of the QTL in our study overlapped known tillering genes or mutants. The Alqudah et al. study (2016) identified tillering QTL that mapped near the low tillering gene *CUL4* (3H, 137.74), but they did not identify other QTL overlapping known tillering genes. The 3H-135 QTL in our study mapped near *CUL4*; however, it is an unlikely candidate gene because LD decays below background levels between 3H-135 and *CUL4* at 137.71 cM (Figure S8). The nearest gene to 3H-135 (HORVU3Hr1G103960, 135.39 cM) encodes an epoxide hydrolase that is more highly expressed in developing tillers than any other tissues based on expression data from Barlex (barlex.barleysequence.org). Another potential candidate gene in this region is homologous to *PLASTOCHRON3*/*GOLIATH* (HORVU3Hr1G104570, 135.39 cM), which encodes a glutamate carboxypeptidase that regulates plastochron length in rice (Kawakatsu *et al.* 2009 p. 3). A plastochron is the time interval between formation of successive leaf primordia (McMaster 2005), and *PLASTOCHRON* mutants are characterized by an increased rate of leaf development. Leaves develop in a phytomer unit that also contains an axillary bud (McMaster 2005), so reduced plastochron (faster leaf development) could also result in higher tiller number under favorable environmental conditions. For example, mutations in *MND4/6*, a gene homologous to rice *PLASTOCHRON1*, causes a high tillering phenotype (Mascher *et al.* 2014).

The low tillering mutants *cul2*, *cul4*, *als*, and *lnt1* are deficient in axillary meristem initiation and maintenance and produce few, if any primary axillary buds (AXB) and no secondary AXB (Babb and Muehlbauer 2003 p. 2; Dabbert *et al.* 2009, 2010; Tavakol *et al.* 2015). Primary AXB form in leaf axils of the main shoot, and secondary AXB form in leaf axils of tillers that develop from primary AXB. Natural variation in primary AXB number has not been assessed, but it is possible that variance in tiller number is influenced more by genes regulating initiation of higher level (secondary, tertiary, etc.) AXB and outgrowth of tillers. Genomewide association studies on the number of secondary and tertiary AXB and outgrowth of tillers, could be a useful way to identify new natural genetic variation for tiller development in barley.

## CONCLUSIONS

Tillering is a complex trait influenced by environment, other traits, and many small effect loci. Based on results of this study it appears that plants utilize resources and make more grain bearing spikes when conditions are favorable, without sacrificing other components of yield, like seed number or weight. In addition, our results and other studies indicate that genetic variation associated with days to heading and spike row-type consistently influences tiller number across different environments. However, identifying genetic variation associated with tiller number in different environments will be essential for gaining a full understanding of the genetic control of tiller development and may be useful for identifying variation suited for adaptation to specific environments.

## ACKNOWLEDGEMENTS

The authors would like to thank Professors Rex Bernardo and Candice Hirsch for assistance with analyses and technicians Shane Heinen, Edward Schiefelbein, and Guillermo Velasquez for assistance with planting. We would also like to thank former lab members: Maria Muñoz-Amatriaín, for providing information about the USDA NSGCC, and Liana Nice, for helping with field design and analysis. We would like to acknowledge the Minnesota Supercomputing Institute (MSI) at the University of Minnesota (UMN) for computing resources (https://www.msi.umn.edu). Finally, special thanks to former undergraduate students who assisted with plant phenotyping and other field work: Allison Shaw, Adam Schrankler, Calandra Sagarsky, Praloy Carlson, Raone Soares Biancardi, Lothi Yamat, Jesus Gonzalez Langa, William Corcoran, Jennifer Nguyen, Amos Kidandaire, Sonya Yermishkin, and Colin Finnegan. This work was financially supported by the USDA-NIFA Triticeae Coordinated Agriculture Project (TCAP, Grant no. 2011-68002-30029).

